# Differential variant calling in mutants from diverse genetic backgrounds: A case study in the nematode *Pristionchus pacificus*

**DOI:** 10.1101/138479

**Authors:** Christian Rödelsperger, Eduardo Moreno

**Affiliations:** Department for Integrative Evolutionary Biology, Max Planck Institute for Developmental Biology, Spemannstr. 35, 762076, Tübingen, Germany

**Keywords:** whole genome sequencing, variant calling, reference bias, assembly error, samtools

## Abstract

Genome sequencing of mutants is one of the most widely used techniques to identify genes that control traits of interest including human diseases. Traditionally, variants are called against reference genomes and various filtering techniques are applied to reduce the number of candidate mutations. However, if the genetic background of the mutant is different from the reference genome, the number of background variants may exceed the number of true mutations by several orders of magnitude resulting in candidate lists that cannot be effectively reduced.

We introduce the problem of differential variant calling in mutants from diverse genetic backgrounds. In the example of a mutant strain of the nematode *Pristionchus pacificus*, where the genetic background is not identical to the reference genome (≈1% genome-wide divergence), we show that simple intersection filtering does not effectively reduce the list of candidate mutations due to the combined effect of high number of background mutations, missing coverage in the wildtype sample, and problematic regions in the genome assembly. Although restriction to sites with coverage in mutant and wildtype sample greatly reduced the number of candidate sites, we further improved this candidate set by implementing a customized variant calling procedure. This takes the mutant sample and the control sample of same genetic background as input and calls variants exhibiting strong discriminative signals across the two samples. Intersecting the candidate mutations with an interval identfied from mapping by RAD-seq revealed a likely splice-site mutation in the *P. pacificus* dpy-1 gene, which has been previously shown to cause the associated morphological phenotype.

Our study shows that combined analysis of mutant and wildtype samples drastically increases the potential to find true mutations. We hope that these results may be helpful for other model systems, where the identification of candidate mutants is complicated by assembly errors, lab-derived mutations, or different genetic backgrounds.

## Introduction

The identification of disease-causing mutations builds an entrypoint to improved diagnosis, understanding disease mechanisms, and ultimately to develop therapies. The advent of next generation sequencing in combination with targeted DNA capturing techniques had tremendous impact on our potential to identify disease-causing mutations. In addition to studies in biomedical contexts [1, 2], evolutionary studies in multiple species with the goal to understand the genetic basis of various traits have made extensive use of forward genetic screens [3, 4, 5, 6, 7]. Typically, the whole genome or a captured fraction of DNA from individuals carrying a mutation is sequenced on one of various next generation sequencing platforms resulting in millions of short reads that are aligned against a reference genome. From the pileup information of bases, aligned to a particular genomic position, differences with respect to the reference sequence are detected. The resulting variants are filtered based on pedigree information [8], common polymorphisms, and disease-causing potential [9] to distill hopefully a small set of candidate mutations. While the low genetic diversity (*≈*1 variant per kilobase) in humans limits the total number of variants and allows for efficient filtering based on common polymorphisms [10], in many other species including *Drosophila melanogaster* and some nematodes, genetic diversity is at least 10 times higher and variant filtering techniques become more challenging. For example, if sequencing of a single exome yields roughly 10-30 thousand variants of which roughly 90% can be filtered out based on polymorphism data [8] and further 70% can be filtered out because they are either intronic or synonymous [2], this results in a set of 300-900 candidate variants that can be further reduced if data from relatives exist. The final candidate set of mutations can be manually checked in visualization programs [11] for alignment artifacts. However, if genetic diversity is ten times higher than in humans, then manually investigating ten times more candidate variants is not an option.

The nematode *Pristionchus pacificus* is one example of a more diverse model organism. Our lab has established *P. pacificus* as a satellite model organsim to *Caenorhabditis elegans* for comparative studies of development [12, 5], ecology [13, 14], and population genomics [15, 16]. Based on world-wild field trips to sample more nematode strains, we have recently discovered one *P. pacificus* strain, RSB001, on a high altitude location on la Reunion Island in the Indian Ocean that shows a so called bordering behaviour (Fig. 1a,b) that has also been observed in *C. elegans* [17] but is not controlled by the same gene [18]. Whole genome resequencing data has shown that this and other strains of the same genetic lineage are roughly 1% divergent from the standard *P. pacificus* reference strain PS312 (isolated in California, USA) [15, 19]. To facilitate a forward genetic screen, for the loss of bordering behaviour in this strain, we first had to generate a mutant line exhibitting a morphological phenotype (e.g. dumpy, Fig. 1c,d). This is necessary in order to screen for successful backcrosses and consequently to reduce the number of background mutations that were induced by chemical mutagenesis. The problem is that *P. pacificus* as well as *C. elegans* are hermaphroditic nematodes that can self-fertilize with males appearing spontaneously at low rate. This facilitates genetic crosses between different strains, but these are complicated as even unsuccessful crossses can lead to progeny due to self-fertilization of hermaphroditic animals. Thus, to discriminate self-offspring from cross-progeny, males carrying the chemically induced mutation resulting in a loss of bordering behaviour need to be crossed with dumpy hermaphrodites so that a successful cross can be detected by the fact that all cross-progeny will be heterozygous at the dumpy locus and display the wildtype body morphology.

**Figure 1:**
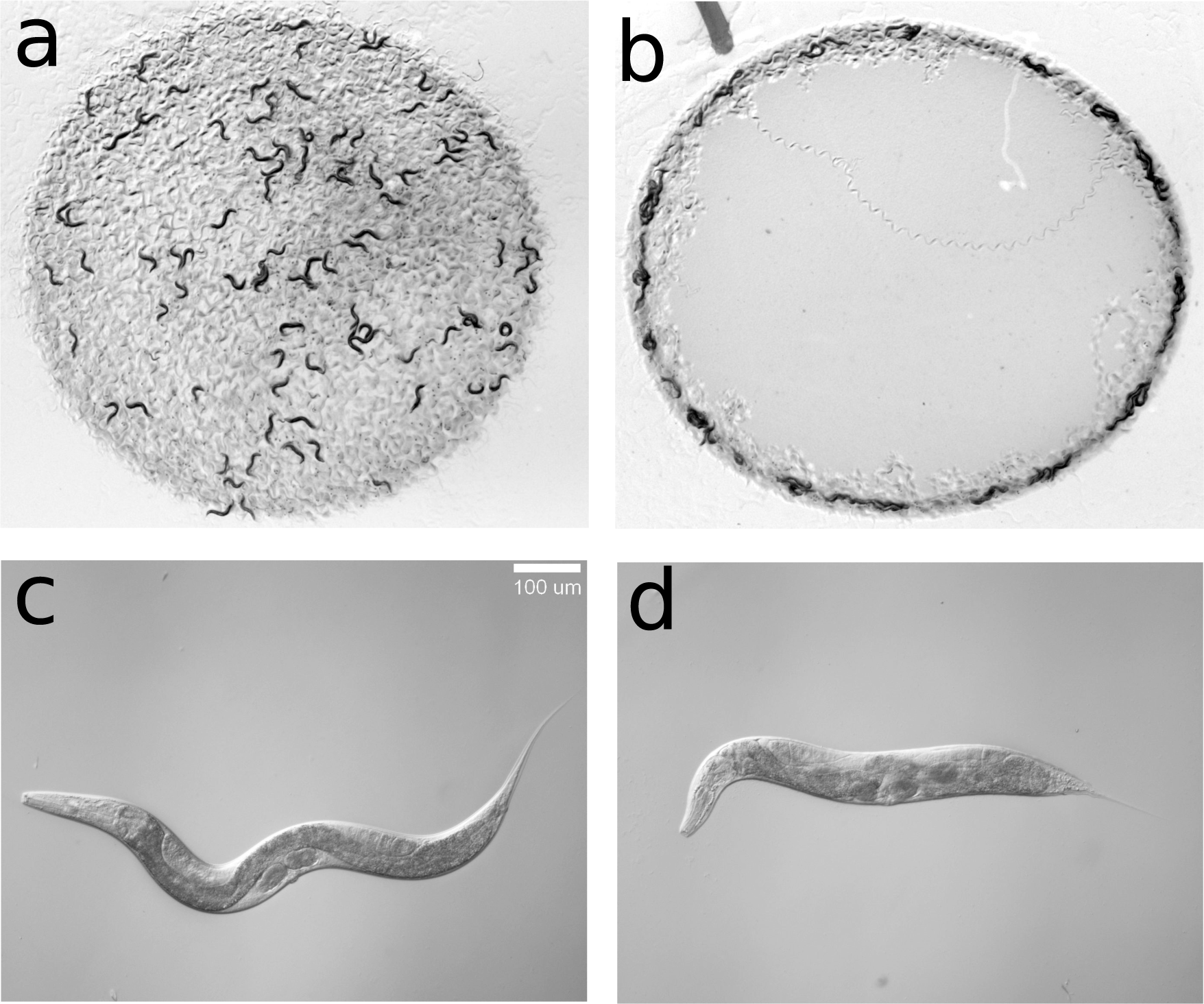
**Bordering bahaviour and dumpy phenotype in *P. pacificus* a)**The reference *P. pacificus* strain PS312 has no preferred localization on plates seeded with *Escherichia coli* OP50. **b)** The strain RSB001 shows a so called bordering behaviour, which has also been observed in *C. elegans* but has a different genetic basis [18]. **c)** A single wildtype worm of *P. pacificus* **d)** A single mutant worm displaying the dumpy phenotype. Morphological markers such as dumpy are frequently used in *C. elegans* and *P. pacificus* to distinguish self from cross-progeny.

In this study we report the generation of a dumpy mutant in the *P. pacificus* strain RSB001 and we identify its molecular identity by means of whole-genome sequencing combined with Restriction enzyme associated DNA tag sequencing (RAD-seq). We found that the analysis of whole genome sequencing data is complicated by the high degree of divergence from the PS312 strain that has been used to assemble the *P. pacificus* reference genome [20] and that simply substracting common variants with the resequencing data of the RSB001 wildtype strain is not practical to define a high-quality set of candidate mutations. We propose that a heuristic variant calling procedure simultaenously taking into account the mutant and the wildtype sample is enough to define more reliable sets of candidate mutations.

## Results

### Whole genome sequencing of a divergent strain

We generated a dumpy mutant in the genetic background of RSB001 (see *Methods)* and resequenced genomic DNA of the mutant (≈ 20× coverage) on a multiplexed run of an Illumina HiSeq 2000 sequencer. The wildtype strain had already been sequenced (≈ 31× coverage) as part of a previous population genomic study [19]. After alignment to the reference genome (153Mb assembled sequence, 173Mb including gaps), we called 1,405,325 and 1,398,744 single nucleotide variants in the mutant and wildtype sample, respectively (Fig. 2a). Removing the intersection between both sets from the mutant sample reduced the candidate set to 68,537 (5%). We selected 100 random sites and manually classified these based on visual inspection of the alignments in the Integrative Genomics Viewer [11] (Fig. 2b). We judged all100 sites as not being different between the two samples. In most cases these candidate sites were not called in the wildtype sample due to little or no coverage, or were due to ambiguous bases that were called homozygous in one sample but heteroygous in the other (Fig. 2b). Based on the observation that in all the cases, where the wildtype sample had little coverage, the alignment of the wildtype sample showed the same base as in the mutant sample, we tend to conclude that also the cases with no coverage in the wildtype sample have most likely the same genotype as the mutant sample. However, since we did not experimentally genotype any of the variants observed in the mutant but not the wildtype, there might still be the chance that some of these sites are truly different between the two samples and that the differences in coverage are due to complex mutations or indels resulting in alignment problems and lack of sequencing coverage. Ambiguous signals in the alignments can be attributed to duplications and assembly problems [15] but also to heterozygosity. As we are looking for a recessive mutation, sites with ambiguous signals can be easily filtered out from the candidate mutant set. This analysis showed that calling variants against the reference, which has worked multiple times effectively when the genetic background of mutant is the same as the reference genome [21, 4, 5, 7], is inappropriate when the genetic background is different. The main problem is that coverage fluctuations can be neglected if the genetic background is the same as the reference, but they drastically inflate the number of candidate mutations in diverse genetic backgrounds (Fig. 2b). The problem of missing coverage prohibits also the approach to change the reference to make it more similar to the strain of interest. Finally, we would like to point out the possibility, that in the worst case, the true mutation could consist in the reversal of the original RSB001 genotype to the genotype of the reference genome, indicating that also the set of 61,956 variant calls that is specific to the wildtype sample has to be considered.

**Figure 2:**
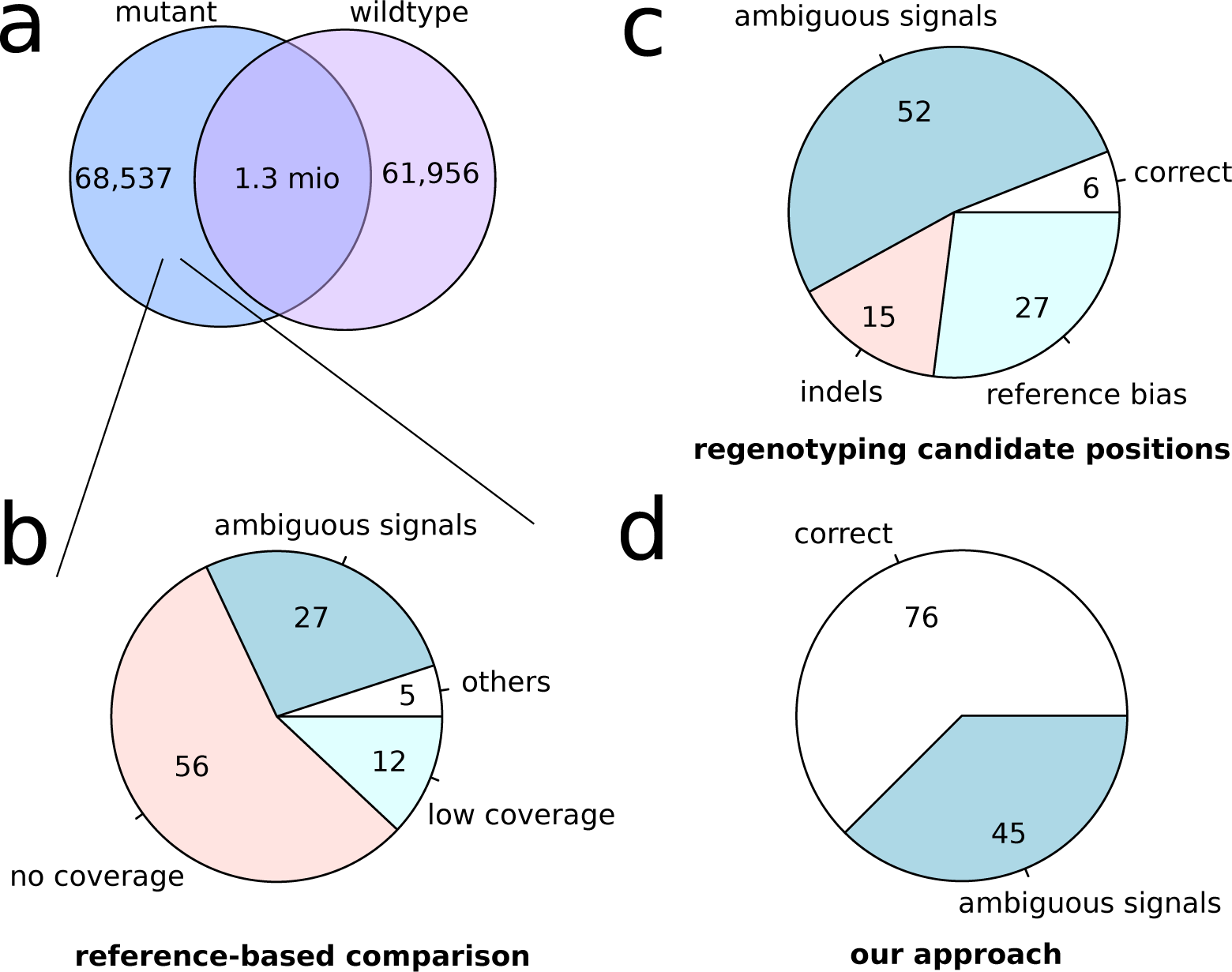
**Variant calling against the reference and intersection filtering** A) Roughly 1.4 million variants wered called against the reference genome for the mutant and wildtype sample. Filtering out all common variants between both samples, reduced the set of candidate mutations to 68,537 (5%). B) Visual inspection of alignments around 100 candidate mutations showed a variant was only called in the mutant sample because of insufficient coverage in the wildtype sample. C) Candidate positions were regenotyped in the mutant sample and 100 randomly selected differential calls with sufficient coverage in both samples were manually inspected and classified in IGV. D) Differential variant calling based on simulataneous analysis of both samples resulted in 121 differential calls.

### Explicit genotype calling at candidate positions

As most of the initial candidate sites are due to low or missing coverage in the wildtype sample, we tested, whether this can be corrected by explicitly calling the genotype at candidate positions in the wildtype sample and only compare those positions that have sufficient coverage but still support a differential variant call between the two samples. Genotyping by a combination of samtools and bcftools removed all but seven positions of the previously 100 cases that were classified as false differential calls (Fig. 2b). The exceptions mostly included positions with ambiguous bases in combination with reads with low mapping quality. To quantify, to what extent, regenotyping helps to reduce the candidate set, we counted how many positions are considered as different between the two samples and have sufficient coverage and are called as homozygous in both samples. This procedure reduced the candidate set to 4,773 (7%) of the initial 68,537 variants. We again chose 100 random positions and investigated the alignments in IGV. The largest number of cases (*N* = 52) still showed ambiguous signals in at least one of the two samples, often combined with low quality mapped reads (Fig. 2c). Further we identified regions with indels [22] that cause discordant variant calls between the samples, but also positions which we classified as “reference bias”. This means that if alignment quality is poor and read coverage low, the variant caller seems to have a tendency to call the reference base. Finally, we found six cases that looked to us as if they represented true differences. At these sites, close to 100% of reads showed one allele in the mutant sample and the other allele in the wildtype sample.

### Strategy for differential variant calling

Altogether, these findings show that without correction for sufficient coverage in both samples, naive analyses of mutants from diverse genetic backgrounds may generate orders of magnitude higher numbers of candidate mutations than there are true differences between the samples. In addition, resequencing of the reference strain PS312 showed roughly 30,000 variants [21, 5, 15] that are probably a result of assembly problems and/or lab-derived mutations since the sequencing of the reference strain [20]. To distill a hiqh quality set of differential variant calls, we therefore decided to implement our own approach, rather than optimizing all the different parameters of the existing variant callers. We decided to analyse both samples simultaenously by parsing the pileup information, which contains all aligned bases at a given position, and employing frequency and coverage thresholds to define candidate sets of differential variants. As the main problem consisted in too high numbers of candidate sites as they were generated by the above mentioned approaches, we aimed to find a very conservative set of high quality variants even if this meant that we might miss some true differences between the strains. Once, this approach has been shown to be successful, the applied thresholds can be lowered in order to find a more complete variant set. We used the candidate variants, that were specific to either data set in Figure 2a, and generated pileup files for 130,493 positions in the mutant and wildtype sample. We then implemented a program, which parses the pileup files, and reported all differential variants that suffice the given coverage (≥ 5 ×) and allele frequencey thresholds (≥ 0.9 in the mutant and < 0.1 in the wildtype sample). This yielded 120 calls which were then manually investigated on IGV. Despite the fact, that we see substantial signal of ambiguous base calls, which could be explained by true heterozygosity, duplicated regions or assembly problems, we classified 76 candidates as correct differential variant calls. Relative to the initial candidate set of 130,493 positions, this represents a lOOO × reduction in the number of candidate mutations, but since we do not know, how high the number false negatives, we cannot make any statement whether 120 is the true number of differences between the two samples. Changing the minimal coverage threshold yield 527 positions with a minimal coverage of two and 1419 positions with a minimal coverage of one, which have to be further checked in case, that the initial candidate set of 75, does not contain the causative mutation.

### Identification of the causative gene

The application of our differential variant calling approach yielded for the first time a set of candidate variants that can be further checked f or functional impact. However, since whole-genome sequencing alone is unlikely to identify a single mutation, we generated some mapping data from crosses between the mutant strain and another strain RSC011 [19]. We scored the morphological phenotype in the offspring and genotyped 57 lines by means of RAD-seq [23]. Based on 201 markers, we found the strongest association on chromosome III (Contig2:1113085, *P <* 10*−*7, Fisher’s excact test, Fig. 3a). While out of the 76 correct variants (Fig. 2D), 10 are predicted to have a functional impact on the protein [21], only one of them was located on the same Contig. At position Contig2:994951 (Hybrid1 assembly), immediately next to the end of the nearest exon, a cytosine was changed into a thymine resulting in a predicted splice-site mutation (Fig. 3b). The gene is the *P. pacificus* ortholog of *C. elegans* dpy-1, which had the same phenotype in *C. elegans* and knockout by the CRISPR/Cas9 system of exactly this gene has shown, that it also has the same phenotype in *P. pacificus* [23]. We therefore conclude that the identified splice-site mutation is the causative mutation resulting in the observed morphological phenotype.

**Figure 3:**
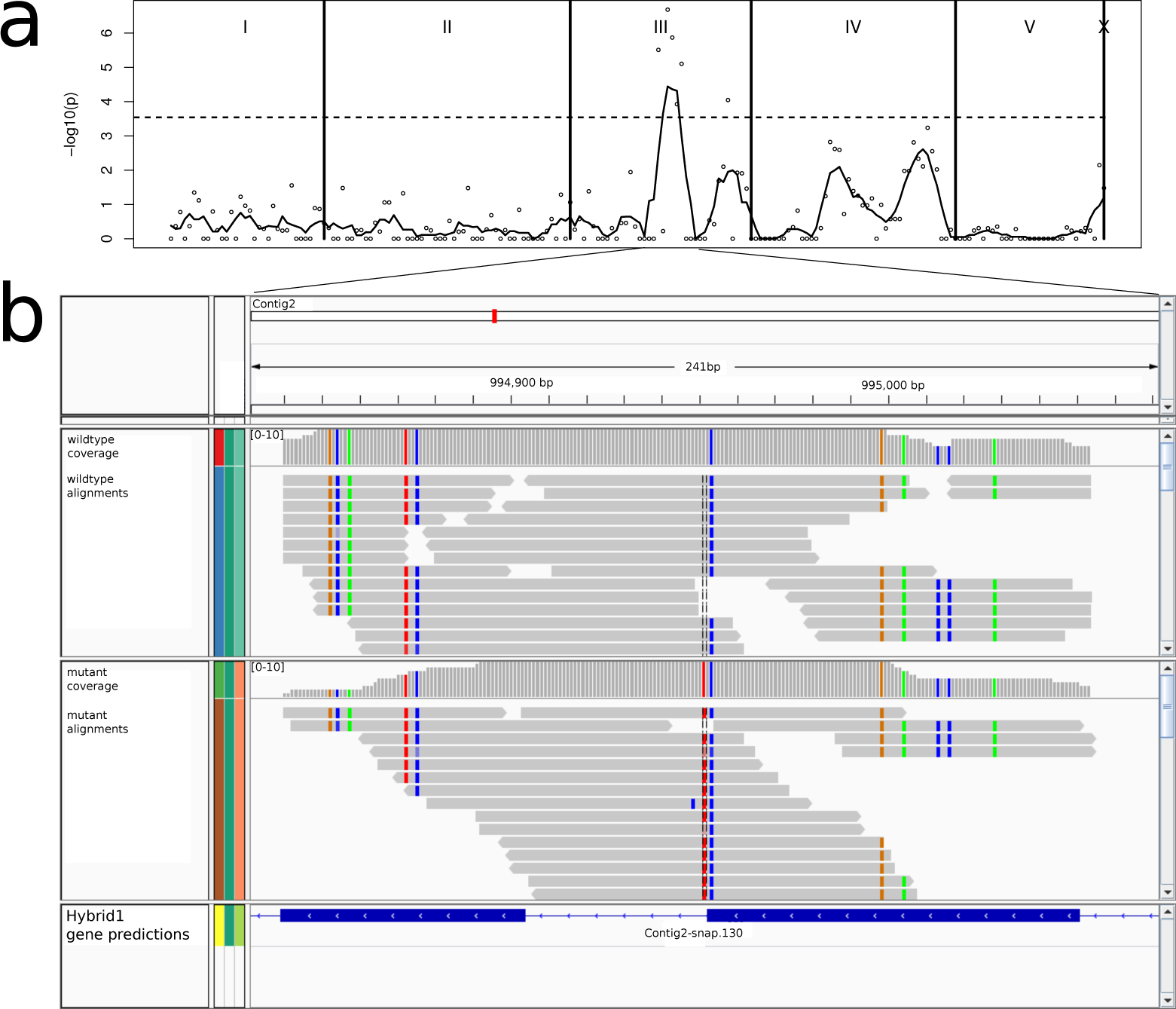
**Identification of the causative gene for the dumpy phenotype a)** The plot shows on the x-axis the relative position of 201 markers, genotyped in 57 mapping lines. The y-axis shows the significance of association between genotype and phenotype. The dashed lines indicates the Bonferroni corrected p-value cutoff. Please note that the lack of X-chromosomal markers is not relevant for this study, since the trait is recessive and hemizygous F1 males do not exhibit a dumpy phenotype (see *Methods*). **b)** One out of ten mutations with predicted functional impact, was located roughly 120kb away from the most significant marker. The mutation was located in a splice site of the *P. pacificus* ortholog of *C. elegans* dpy-1.

## Discussion

The main purpose of this study is to explicitly describe the problem of calling differential variants in diverse genetic backgrounds. Although widely used software suites such as samtools [24] FreeBayes [25], or GATK [26] can in theory be employed to obtain similar results, to our knowledge, there is no study that explicitly deals with this subject and variant callers do not offer the option to specify a control sample as additional input file. The problem of differential variant calling is related to the identification of tumor specific mutations relative to healthy tissues [27]. However, there are two important differences, the first being that genetic background is more or less the same as for the human reference, and the second being that it is never clear how pure the healthy and tumor samples are. Apart from the identification of causative mutations in forward genetic screens, differential variant calling can be employed to identify reliable markers for quantitative trait loci studies where both parental lines differ from the reference genome. The alternative to assemble a genome *de novo* for each strain that is subject to genetic studies is probably doable but not really cost effective. Even for variant calling in samples from the same genetic background as the reference, differential variant calling relative to the wildtype sample may be useful as an alternative strategy to directly remove false variant calls introduced by errors in the genome assembly [28].

In multiple previous studies, we have demonstrated that calling variants against the reference is sufficient to identify causative mutations in the same genetic background as the reference [21, 14, 5, 7]. Here we show that it produces an unmanagable number of candidate mutations if the genetic background differs from the reference. This is because the shear number of variants, that are due to the different genetic background but is not covered in the wildtype sample, drastically inflates the number of candidate muations. While this can also not be solved by changing the reference sequence to be more similar to the strain of interest, just correcting for sites that are effectively covered in both samples already greatly reduced the number of candidate variants. Further implementation of an intuitive method for differential variant calling in order to reduce the amount of ambiguous candidates allowed us to identify the causative gene for the *P. pacificus* dumpy mutant line.

Finally, we would like to emphasize that classifying candidate mutations based on visual inspection of the alignments is a fast way to get a feeling of the quality of variant calls, but also simultaneously generates a test set that can be used to re-adjust the employed frequency and coverage cutoffs. These may differ depending on the rate of sequencing errors and quality filtering methods (e.g. quality based read trimming, realignment, removal of duplicate reads). Similarly, frequency differences in regions of high coverage are inherently more significant than in regions of mean coverage, yet the fact that a region has high coverage is already a good indicator for unreliable variant calls as they might be affected by heavily amplified, repetitive, or duplicated sequences. On hindsight, this empirical approach is subjective and might heavily depend on the ability of the user to recognize artifacts such as false variant calls that are due to misaligned indels [22].

## Conclusions

Our study has shown that the genetic background of the reference genome has a huge impact on the methods that need to be employed for variant calling. We have proposed differential variant calling strategies that simultaneously take into account the mutant and wildtype sample, as a means to reduce the effect of a diverse genetic backgrounds and we have used such an approach to identify the causative gene for a dumpy phenotype in the nematode *P. pacificus*. We hope that this example will help other studies aiming for mechanistic insights into the genetic basis of various traits in non-classical model organisms.

## Methods

### Worm culture and mutagenesis

Strains were maintained at 20 °C using standard methods [29]. A dumpy mutant in the genetic background of RSB001 was generated by means of a standard mutagenesis protocol [30]. This mutant was backcrossed twice in order to reduce the number of background mutations in the dumpy line. To test for dominance we analyzed the progeny of a cross between dumpy hermaphrodites and RSB001 wildtype males. The appearance of wildtype hermaphrodites among the hybrid F1 nematodes indicated that the dumpy phenotype is fully recessive. On the other hand, the hybrid F1 males produced by the cross did not show the dumpy phenotype. Since males only posses one copy of the X chromosome, which they inherit from the mother, this experiment indicated that the locus responsible for the dumpy phenotype was not located on the X chromosome.

### Whole genome and RAD sequencing

A pellet of dumpy mutant nematodes for genomic DNA extraction was obtained as described in Witte et al [23]. The genomic DNA was extracted using the GenElute Mammalian Genomic DNA Miniprep Kit from Sigma-Aldrich (Sigma-Aldrich Chemie GmbH, Taufkirchen, Germany) and quantified with the Qubit dsDNA HS Assay Kit from Thermo Fisher Scientific (Life Technologies GmbH, Darmstadt, Germany). A genomic library was generated using the TruSeq Nano DNA Library Preparation Kit from Illumina (Illumina Inc., California, United States) following the kit protocol. The library was validated on an Agilent Bioanalyzer DNA 1,000 chip (Agilent Technologies GmbH, Waldbronn, Germany) before sequencing on an Illumina HiSeq 2000 sequencer.

Recombinant lines were constructed by mating wildtype males of the RSC011 strain with hermaphrodites of the newly generated dumpy line in the RSB001 genetic background. From wildtype hybrid F1 hermaphrodites, 57 F2 hermaphrodites were isolated after screening for the dumpy phenotype in order to produce a set of mapping lines containing approximately the same number of wildtype and mutant lines. Finally these lines were genotyped by means of RAD-seq as indicated in Witte et al [23].

### Alignment and variant calling against the reference

We aligned raw paired-end reads (2 × 101nt) of the RSB001 wildtype strain (26.6 × 106 read pairs) and the corresponding dumpy mutant (17.3×106 read pairs) with the help of the BWA program (default options, version 0.7.5a-r405) [31] against the *P. pacificus* reference genome (version Hybrid1, 153Mb assembled sequence, downloaded from pristionchus.org). Initial variant calls against the reference genome were generated by a combination of samtools mpileup (options: -uf, version 0.1.19-96b5f2294a), bcftools view (options: -vcg), and vcfutils.pl varFilter (options: -D1000 -w0) [24]. The exact program call is:

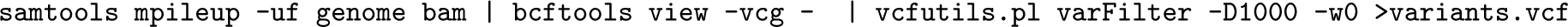

where genome denotes the fasta file with the genome assembly, bam denotes the alignment file, and variants denotes the output variant file. Variants with a quality score below 20, or only supported by a single read (DP*<* 2), indels, and apparent heterozygous sites (FQ*>* 0) in the resulting vcf file were removed. We then filtered the mutant file based on variants that were also called in the wildtype sample (identity was determined by exact position and genotype). We then randomly chose 100 sites that were manually classified as true and false differential calls based on visualization of alignments in the Integrative Genomics Viewer (version 2.2.5) [11]. For visualization in IGV we typically filtered out reads with a mapping quality below 20, but if we could not explain the observed variant calls, we enabled the display of all reads.

### Explicit genotype calling at candidate positions

We genotyped 68,537 positions (Fig. 2a) in the mutant and wildtype sample by executing samtools and bcftools

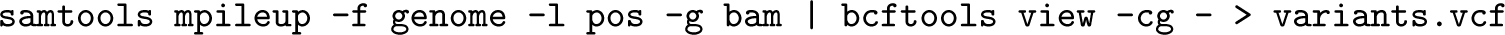

where pos denotes the position file, genome denotes the fasta file with the genome assembly, bam denotes the alignment file, and variants denotes the output variant file. The resulting variant files where filtered for quality ≥ 20, coverage ≥ 2, homozygosity (*FQ* < 0). After further removing sites that are labeled as indels, the mutant and wildtype variant files were screened for positions with calls in both files but differing genotypes (Fig. 2c).

### Strategy for differential variant calling

We first extracted pileup information for the union of all variable sites from the mutant and wildtype sample with the help of the samtools mpileup command (options: -A -q 2):

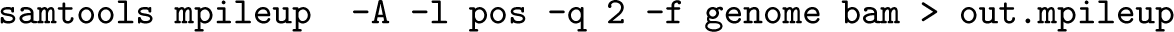

where pos denotes the position file tab separated format (Contig Pos), genome denotes the fasta file with the genome assembly, bam denotes the alignment file, and out.mpileup denotes the output pileup file. We then wrote a perl script, which parses the pileup information for mutant and wildtype sample and calls candidate sites based on thresholds for the frequency (*f* = 0.9, default value) of alleles in both samples and a minimal coverage (*c* = 2, default value). A frequency threshold of *f* = 0.9 indicates that the major allele at a given position in the mutant has to have a frequency ≥ 0.9 whereas in the wildtype sample the frequency of the same allele should be ≤ 1 − *f* = 0.1.

### Analysis of RAD sequencing data

RAD-seq data was analysed as described previously in Witte et al. (2015) [23]. In short, reads were aligned by the BWA aligner (version 0.6.1-r104) against the *P. pacificus* Hybrid1 assembly. Variable positions between RSB001 and RSC011 were taken from the data set by McGaughran et al. [19] and genotyped using samtools. The significance of the association between genotype and phenotype was assessed by performing a Fisher’s exact test.

## Author’s contributions

CR and EM conceived the study. EM carried out the experiments. CR analyzed the data. CR and EM wrote the manuscript. CR and EM have read and approved the final version of the manuscript.

## Acknowledgements

We would like to thank two anonymous reviewers for helpful comments on the manuscript.

## Ethics statement

This study does not involve research on humans or human material and also not on animals according to the german animal protection legislation. Therefore no ethical approval is needed.

## Availability of Data

Raw reads for the wildtype RSB001 strain have been submitted as part of the study by McGaughran et al. (2016) [19] to the European Nucleotide archive (Study accession PRJEB13695, Run accession ERR1390229). Resequencing data for the EMS mutagenized dumpy strain has been submitted to the NCBI sequence read archive (Study accession: PRJNA362524, Run accession SRR5187774).

